# The Role of the Ovarian Cancer G -Coupled Receptor (OGR1) in Idiopathic Pulmonary Fibrosis

**DOI:** 10.1101/849117

**Authors:** David J. Nagel, Ryan Clough, Tyler J. Bell, Wei-Yao Ku, Patricia J. Sime, R. M. Kottmann

## Abstract

Idiopathic pulmonary fibrosis (IPF) is a disease characterized by irreversible scarring of the lung that is associated with significant mortality and morbidity. The pathophysiology is incompletely understood but it is well-established that fibroblast to myofibroblast differentiation is a key feature of pulmonary fibrosis. Our lab has established that a reduction in extracellular pH is one of several important pathways responsible for the activation of latent TGF-β in the extracellular space. TGF-β activation further decreases extracellular pH and creates a feed-forward mechanism that stimulates myofibroblast differentiation and activation of additional TGF-β. Given the importance of TGF-β and extracellular acidification to the progression of pulmonary fibrosis, we sought to identify novel mechanisms that are involved in pH-dependent fibrotic signaling. The proton sensing G-Protein Coupled family of receptors are activated in acidic environments, but their role in fibrotic signaling has not been studied. Here we report that the Ovarian Cancer G-Protein Coupled Receptor1 (OGR1 or GPR68), a member of the family of proton sensing G-Protein Coupled Receptors, negatively regulates pro-fibrotic signaling. We demonstrate that OGR1 expression is significantly reduced in lung tissue from patients with IPF and TGF-β decreases OGR1 expression. In fibroblasts, a reduction in expression of OGR1 (OGR knockout lung fibroblasts) and knockdown (OGR siRNA), promotes *in vitro* myofibroblast differentiation. In contrast, OGR1 overexpression inhibits myofibroblast differentiation. Finally, we demonstrate that OGR1 negatively regulates TGF-β stimulation through inhibition of focal adhesion kinase (FAK) phosphorylation. Our results suggest that preserving OGR1 expression may represent a novel therapeutic strategy in pulmonary fibrosis.

## Introduction

Idiopathic pulmonary fibrosis (IPF) is an unrelenting fibrosing interstitial pneumonia with a median survival of approximately 3 years from the time of diagnosis [1]. In 2014, two new medications were approved for the treatment of IPF (Nintedanib and Pirfenidone)[2, 3]. Each of these medications slow the rate of lung function decline but do not reverse fibrosis. At present, the only curative treatment is lung transplantation. Unfortunately, not all patients are eligible for transplantation and post-transplant morbidity can be substantial.

The underlying mechanisms leading to the development of pulmonary fibrosis remain incompletely understood. One well-established pathway involves the differentiation of fibroblasts to myofibroblasts[4]. Myofibroblasts are one of the primary cell types that produce extracellular matrix proteins [5-8]. Transforming growth factor-beta (TGF-β) is the predominant pro-fibrotic cytokine responsible for the induction of myofibroblast differentiation[9-11]. TGF-β is produced as a latent protein that requires activation through separation from the latency associated peptide (LAP)[12]. Activation of TGF-β occurs through several mechanisms including enzymatic degradation of LAP, mechanical strain, interactions with alpha integrins at the cell surface, and changes in pH and temperature[13-17]. We have demonstrated that TGF-β induces fibroblasts to generate excess lactic acid via the induction of lactate dehydrogenase (LDH), which in turn lowers extracellular pH and activates extracellular latent TGF-β[18]. We have shown that LDH is upregulated in the lung tissue of patients with IPF and in the lung tissue of mice treated with bleomycin[18]. Furthermore, we have shown that the pH of lung tissue of mice exposed to bleomycin decreases to an average of 6.7[19]. Thus, we proposed that pH dependent changes in lung tissue may critically regulate the development of pulmonary fibrosis.

We were therefore interested in exploring other pH-dependent processes that may regulate the development of pulmonary fibrosis. One potentially important pathway involves the expression and activation of proton sensing G-protein Coupled Receptors (GPCR). Little is known about the role of the proton-sensing GPCRs in pulmonary fibrosis although they are increasingly recognized for being pathologically dysregulated in malignancy[20-22], heart failure[23], hypertension[24], pulmonary hypertension[25], irritable bowel disease[26-28], and asthma[25]. There are four members of this family of receptors, the Ovarian Cancer G-Protein Coupled Receptor Protein1 (OGR1, GPR68), G-protein Coupled Receptor 4 (GPR4), G2 accumulation (G2A), and T Cell Death-Associated Gene 8 (TDAG8)[29-32]. The members of this family sense extracellular protons through histidine residues located on the extracellular portion of the receptor[30, 33] which then activate downstream G proteins. OGR1 was originally cloned from an ovarian cancer cell line[34] but has subsequently been found to be expressed in the spleen, testis, small intestine, kidney, brain, heart, and lung[29]. Although several endogenous ligands have been proposed (sphingosylphosphorylcholine, galactosylsphingosine, and lysophosphatidylcholine), the role of OGR1-ligand interaction remains largely unknown[35, 36].

OGR1 is essentially inactive at a pH of 7.8 but fully active at a pH of 6.8[30] where it exerts ligand-independent constitutive activity[37]. *In vitro* expression of this receptor has been shown to suppress metastasis in prostate cancer[37], mitigate cell migration in breast cancer [38], and regulate acid-induced apoptosis in endplate chondrocytes[39]. However, in other studies the absence or reduction of OGR1 expression inhibited melanoma tumorigenesis[40] and was protective against inflammation in an IBD mouse model[27]. In contrast the absence of OGR1 has also been associated with an increased sensitization of asthmatic airways to inflammatory stimuli[41]. Thus, the role of OGR1 in human disease is likely highly dependent on the degree of receptor expression, the local environment, and organ or cell specific expression patterns. We sought to determine if OGR1 expression was dysregulated in pulmonary fibrosis and if a reduction in OGR1 expression in lung fibroblasts would modulate myofibroblast differentiation. Our data demonstrate that OGR1 is downregulated in pulmonary fibrosis and that OGR1 over-expression negatively regulates pro-fibrotic signaling in lung fibroblasts. Therefore, we hypothesize that OGR1 represents a novel therapeutic target in pulmonary fibrosis.

## Methods

### Human Lung Biopsy Samples

Lung tissue samples were obtained from patients with and without IPF using existing URMC RSRB approved human subject protocols and the Lung Tissue Research Consortium. All tissue was de-identified and collected in a standard manner according to approved institutional review board protocols.

### Tissue Histology/Immunohistochemistry

Mouse lung tissue was prepared according to a standardized protocol [42]. The left lobe was inflated and fixed with formalin. Sections were cut into 5 μM thick sections. Slides were then treated with 3% hydrogen peroxide after re-hydration and antigen retrieval was performed with hot sodium citrate buffer. Slides were then incubated in 5% normal goat serum in phosphate buffered saline with tween for 1 hour. Primary incubation occurred overnight at 4°C with the following antibodies: αSMA (Sigma-Aldrich, St. Louis, MO) and OGR1 (Exalpha Biological Inc, Shirley, MA). After washing, biotinylated secondary antibodies (Jackson ImmunoResearch, West Grove, PA) were incubated for 1 hour. Next, Streptavidin-HRP (Jackson ImmunoResearch) was incubated for 15 minutes. Slides were then developed with 3,3’ Diaminobenzidine (DAB) (Vector, Burlingame, CA). Slides were counterstained with Hematoxylin (BioCare Medical, Concord, CA) and mounted with a coverslip and VectaMount mounting media (Vector, Burlingame, CA). Slide images were acquired with an Olympus microscope and camera (Olympus, Tokyo, Japan) using identical settings and post-capture processing.

### Primary Fibroblast Culture

Human lung fibroblasts were cultured, under a protocol approved by the Institutional Review Board at the University of Rochester Medical Center, from explanted tissue obtained from either organ donors or those undergoing surgical lung biopsy. Primary mouse lung fibroblasts were cultured from lung tissue from 8-10 week old wild type, C57Bl/6J mice (Jackson Labs, Bar Harbor, ME) and OGR1 global knockout mice (Novartis, Basel Switzerland). All fibroblasts were maintained in Modified Eagle Medium (MEM) supplemented with 10% fetal bovine serum, 1% antibiotic-antimycotic, and 1% L-glutamine. Passages from 3 to 9 were used for all experiments.

OGR over-expression was accomplished using the X-treme Gene HP protocol (Roche, Basel Switzerland) and an OGR plasmid previously described [43]. Briefly, fibroblasts were plated at a density of 7×10^4^ in 12-well tissue culture plates 24 hours prior to transfection. Cells were plated in MEM with serum as above, but plasmid DNA was delivered in serum-free OptiMEM (Gibco, Fisher Scientific, Waltham MA). A ratio of 3:1 (3 µL of HP reagent: 1 µg of DNA) was combined with Opti-MEM in a sterile tube and incubated for 15 minutes at room temperature. The transfection complex was then added to each well in a dropwise manner. After incubation for a minimum of 18 hours, cells were administered TGF-β (1 ng/mL) or DMSO (equal volume to that of TGF-β).

OGR expression was knocked down in human lung fibroblasts with SMARTpool ON-TARGETplus GPR68 siRNA (Dharmacon, Lafayette, CO). Cells were plated at 7×10^4^ in 12 well plates. The X-treme Gene SiRNA reagent was utilized (Roche, Basel Switzerland) in combination with Opti-MEM media per manufacturer instructions. The following morning, Opti-MEM media was aspirated and cells were treated with TGF-β or DMSO in serum-containing MEM media.

Fibroblast stimulation of actively growing mouse and human primary lung fibroblasts was achieved with administration of TGF-β (1 ng/mL). Cells were maintained in MEM as above; just prior to TGF-β administration, culture dishes were washed with sterile phosphate buffered saline. TGF-β or dimethyl sulfoxide (DMSO, control) was combined with MEM media and after thorough mixing, was applied evenly to each appropriate well. Treatment occurred for 24-48 hours and then experiments were performed as outlined in other sections.

### Quantitative Real-time Polymerase Chain Reaction

Total RNA was isolated from the right middle lobe of mice and from pathologic samples from human lung tissue using Trizol (Invitrogen, Carlsbad, CA). Reverse transcription utilized the iScript cDNA synthesis kit (Bio-Rad, Hercules, CA). Real-time PCR reactions, using this cDNA (50 ng per reaction), were performed with SYBR Green (Bio-Rad) and were analyzed with a T100 Thermal Cycler (Bio-Rad). The following sequences were used for primers: housekeeping gene (**18S**) <u>forward</u>: 5’GCTTGCTCGCGCTTCCTTACCT, <u>reverse</u>: 5’TCACTGTACCGGCCGTGCGTA; **OGR1** <u>forward</u>: 5’TGTACCATCGACCATACCATCC, <u>reverse</u>: 5’GGTAGCCGAAGTAGAGGGACA; **Collagen 1a1** <u>forward</u>: 5’CTGCTGGCAAAGATGGAG, <u>reverse</u>:5’ACCAGGAAGACCCTGGAATC; **Collagen 3a1** <u>forward</u>: 5’AAATGGCATCCCAGGAG, <u>reverse</u>: 5’ATCTCGGCCAGGTTCTC; **α-Smooth muscle actin** <u>forward</u>: 5’GTGTTGCCCCTGAAGAGCAT3’, <u>reverse</u>: 5’GCTGGGACATTGAAAGTCTCA3’; **Fibronectin** <u>forward</u>: 5’TTGAAGGAGGATGTTCCCATCT3’, <u>reverse</u>: ACAGACACATATTTGGCATGGTT3’.

### Western Blotting

Western blots were performed using total right lung lobe homogenates from mice and human samples as previously reported[44]. Polyvinylidene difluoride (PVDF) membranes (EMD Millipore, Billerica, MA) were activated in methanol and incubated with 5% milk in PBST for 1 hour followed by incubation with the following primary antibodies: OGR1 (“GPR68” Invitrogen, Carlsbad, CA), beta-tubulin (Abcam, Cambridge, MA), α-SMA (Sigma-Aldrich, St. Louis, MO). Goat anti-rabbit and goat anti-mouse secondary antibodies were purchased from Jackson ImmunoResearch (West Grove, PA). Protein bands were developed with Western Lightning Enhanced Chemiluminescence Substrate (Perkin-Elmer, Waltham, MA) and detected on BioBlot plain X-ray film (LPS, Rochester, NY). Densitometric analysis was performed with Image Studio Lite software (LI-COR, Lincoln, NE).

### Data Analysis

Statistical analysis using two-way ANOVA with Tukey multiple comparison method and unpaired T-tests were performed with Graph Pad Prism version 8 (San Diego, CA). Western blot samples represented a minimum of 6 comparators per groups from at least three replicated experiments. Data are expressed at mean ± standard error mean (SEM). A p value of < 0.05 was considered statistically significant.

## Results

### OGR1 Expression is Down Regulated in Lung Tissue from Patients with Idiopathic Pulmonary Fibrosis

Because the extracellular space is acidified in pulmonary fibrosis[45], we were interested in profiling changes in expression for candidate proteins that are involved in proton-mediated signaling. In healthy patients, OGR1 is highly expressed as detected by western blot analysis (Figure 1A, B). In patients with a clinical diagnosis of IPF, OGR1 expression was significantly lower (Figure 1A, B). Immunohistochemistry staining demonstrated that OGR1 was expressed in airway smooth muscle cells, airway epithelial cells, and macrophages (Figure 1C, top left panel). However, OGR1 expression was significantly downregulated in patients with IPF (Figure 1C top right and bottom panels). Interestingly, OGR1 expression in epithelial cells and vascular smooth muscle cells were similar among healthy and diseased patients. Fibroblast expression of OGR1 was significantly reduced in IPF patients, specifically in fibroblastic foci, a pathologic hallmark of the disease (Figure 1 C, arrows). These results demonstrate that OGR1 expression is down-regulated in a fibrotic.

**Figure 1.**
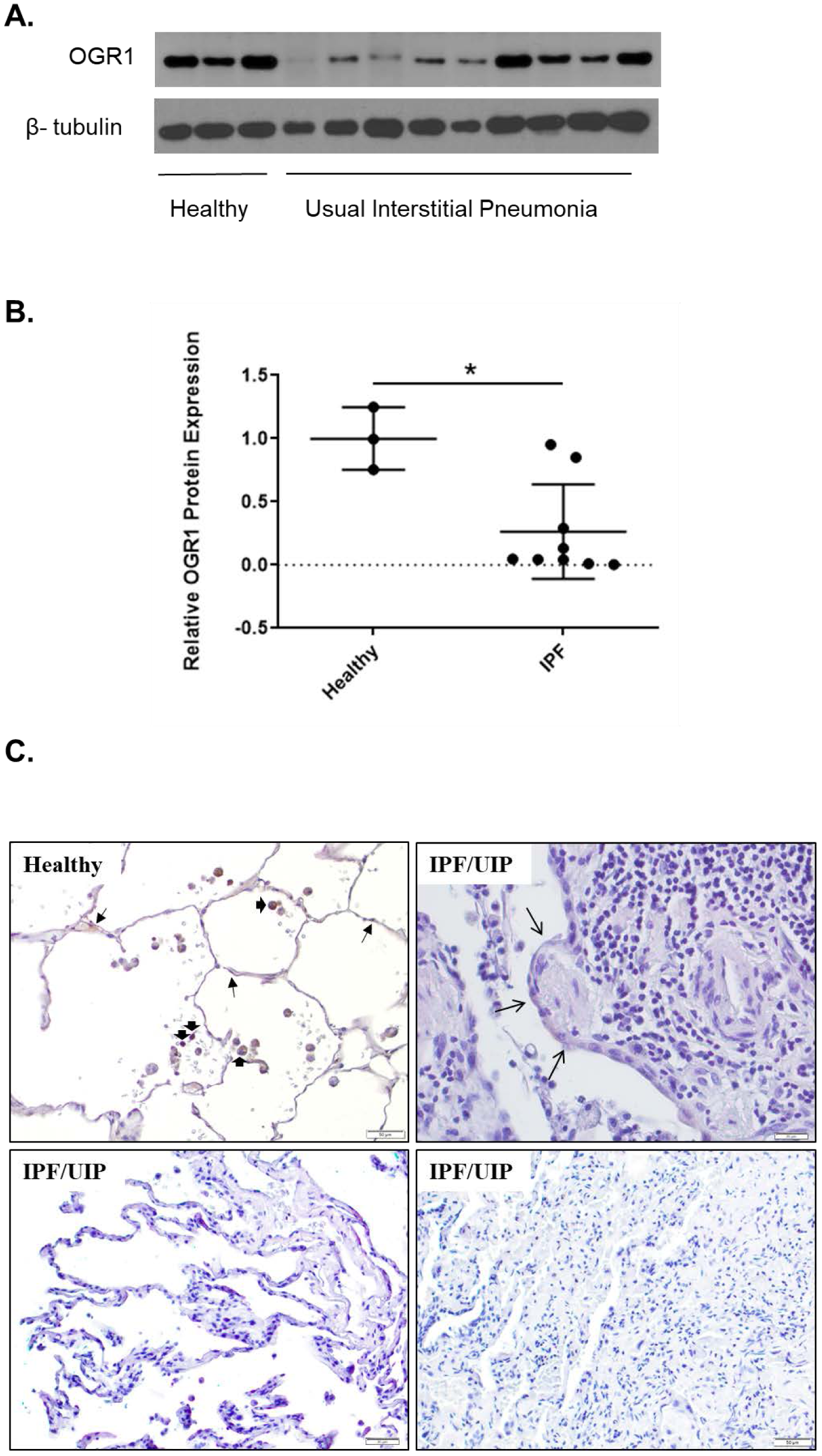
(A) Lung tissues from healthy patients and those with IPF were analyzed for OGR1 protein expression by Western blot. (B) Densitometric analysis was performed and data represents mean OGR1 expression relative to β-tubulin ± SEM. * p = 0.0105. (C) OGR-1 protein expression is down-regulated in pathologic tissue of patients with UIP/IPF. The top left panel represents lung tissue from a healthy donor. There was no staining with Isotype IgG control of the OGR-1 antibody (data not shown). The right upper panel represents a fibroblastic focus (arrows) and demonstrates down-regulation of OGR-1. The bottom panels represent lung tissue from additional patients with UIP/IPF that demonstrate a similar pattern.

### OGR1 Downregulation Induces *in vitro* Matrix Remodeling in HLFs

We hypothesized that reduced OGR1 expression would be associated with a fibrotic phenotype even in the absence of TGF-β stimulation. To assess if downregulating OGR1 effects myofibroblast differentiation, we examined the consequences of using siRNA to decrease OGR1 expression. We observed an approximate 75% reduction in OGR1 after siRNA administration (Figure 2A). Decreasing OGR1 expression led to statistically significant increases in expression of markers of the myofibroblast phenotype including collagen 1A1 (Figure 2B), collagen 3A1 (Figure 2C), fibronectin (Figure 2D), and alpha smooth muscle actin (Figure 2E). All of these proteins have been associated with the extracellular matrix remodeling and myofibroblast differentiation observed in pulmonary fibrosis. This data suggests that OGR1 is an important negative regulator of myofibroblast differentiation.

**Figure 2.**
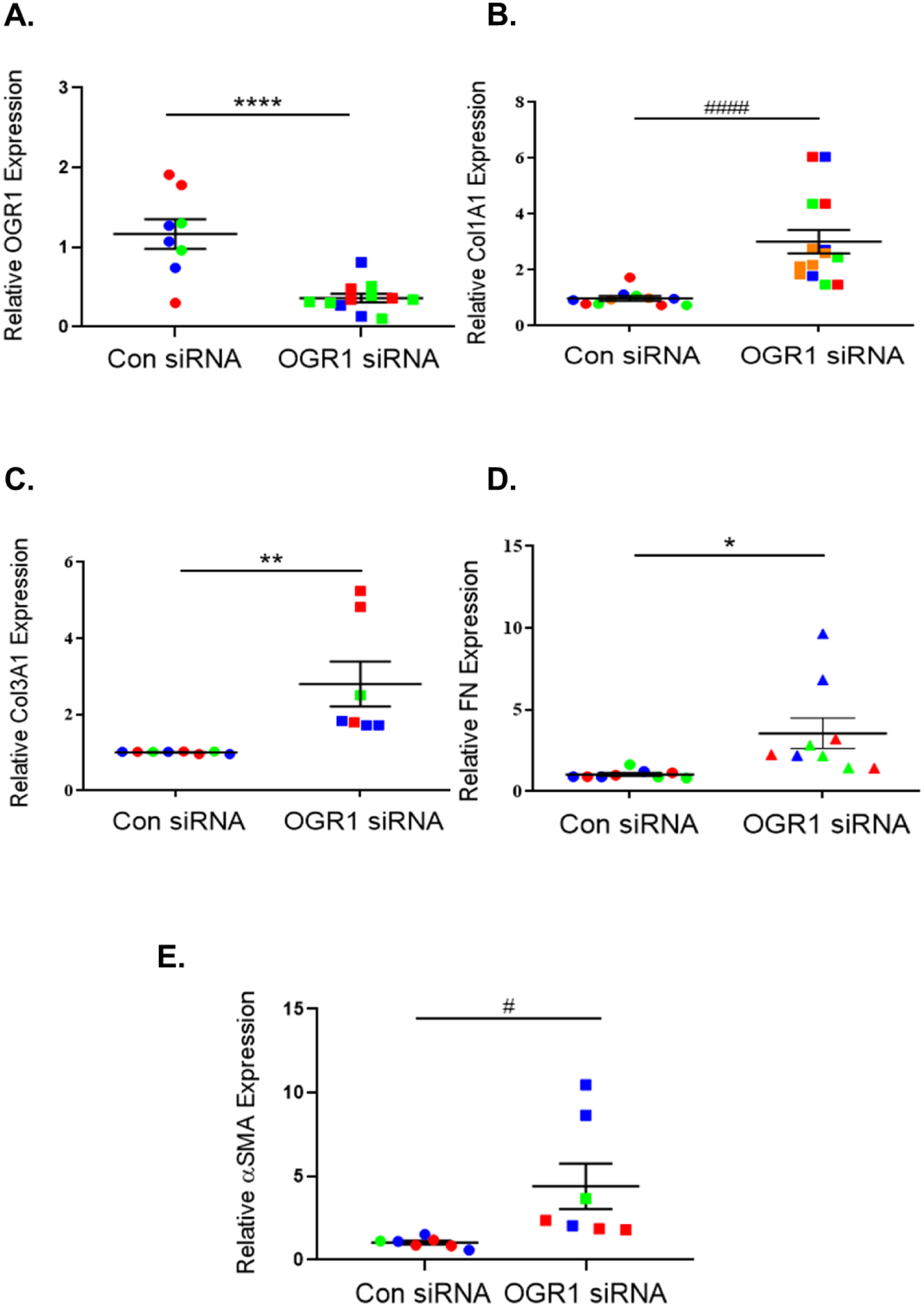
Healthy human lung fibroblasts were treated with control or OGR1 siRNA for 48 hours. The expression of OGR1 (A), collagen 1A1 (B), collagen 3A1 (C), fibronectin (D), and α-SMA (E) were subsequently assessed by qRT-PCR. Results are displayed as candidate mRNA expression relative to 18 S, each set of colors represent a unique cell line isolated from different donors. Data represent mean expression ± SEM (n=3/treatment group, repeated in triplicate in three different cell lines). **** p = 0.0001 (A), #### p = 0.0003 (B), ** p = 0.0058 (C), * p = 0.0165 (D), # p = 0.0303 (E).

### OGR -/- Induces *in vitro* Collagen in MLFs

Because TGF-β is a key cytokine responsible for inducing myofibroblast differentiation[11], and OGR1 expression is down-regulated in myofibroblasts, we wanted to assess whether TGF-β regulates OGR1 expression. Since OGR1 is differentially expressed in IPF lung tissue, we also explored whether TGF-β would have differential effects on OGR1 expression in healthy versus fibrotic fibroblasts. Healthy and fibrotic human lung fibroblasts were treated with TGF-β and change in OGR1 expression was assessed by quantitative real-time PCR and Western blot. Interestingly, baseline OGR1 expression was not different among healthy and fibrotic fibroblasts. However, when healthy fibroblasts were stimulated with TGF-β, there was a significant reduction in OGR1 expression (Figure 3A). Fibrotic fibroblasts demonstrated an even more significant down-regulation of OGR1 after TGF-β administration (Figure 3A). To examine the potential role of OGR1 in myofibroblast inhibition, we next administered TGF-β to wild type and OGR1 knock out mouse lung fibroblasts (MLFs) and examined changes in α-SMA protein expression as a marker of myofibroblast differentiation. In unstimulated states, OGR1 KO MLFs expressed a non-statistically significant increase in α-SMA (Figure 3B). As expected, TGF-β stimulation of wild type MLFs led to a dramatic increase in α-SMA expression (Figure 3 B, C). Importantly, OGR KO MLFs displayed an even more dramatic increase in α-SMA expression (Figure 3B, C). These results demonstrate that the absence of OGR1 sensitizes lung fibroblasts to the pro-fibrotic effects of TGF-β, and therefore suggests that OGR1 negatively regulates TGF-β signaling.

**Figure 3.**
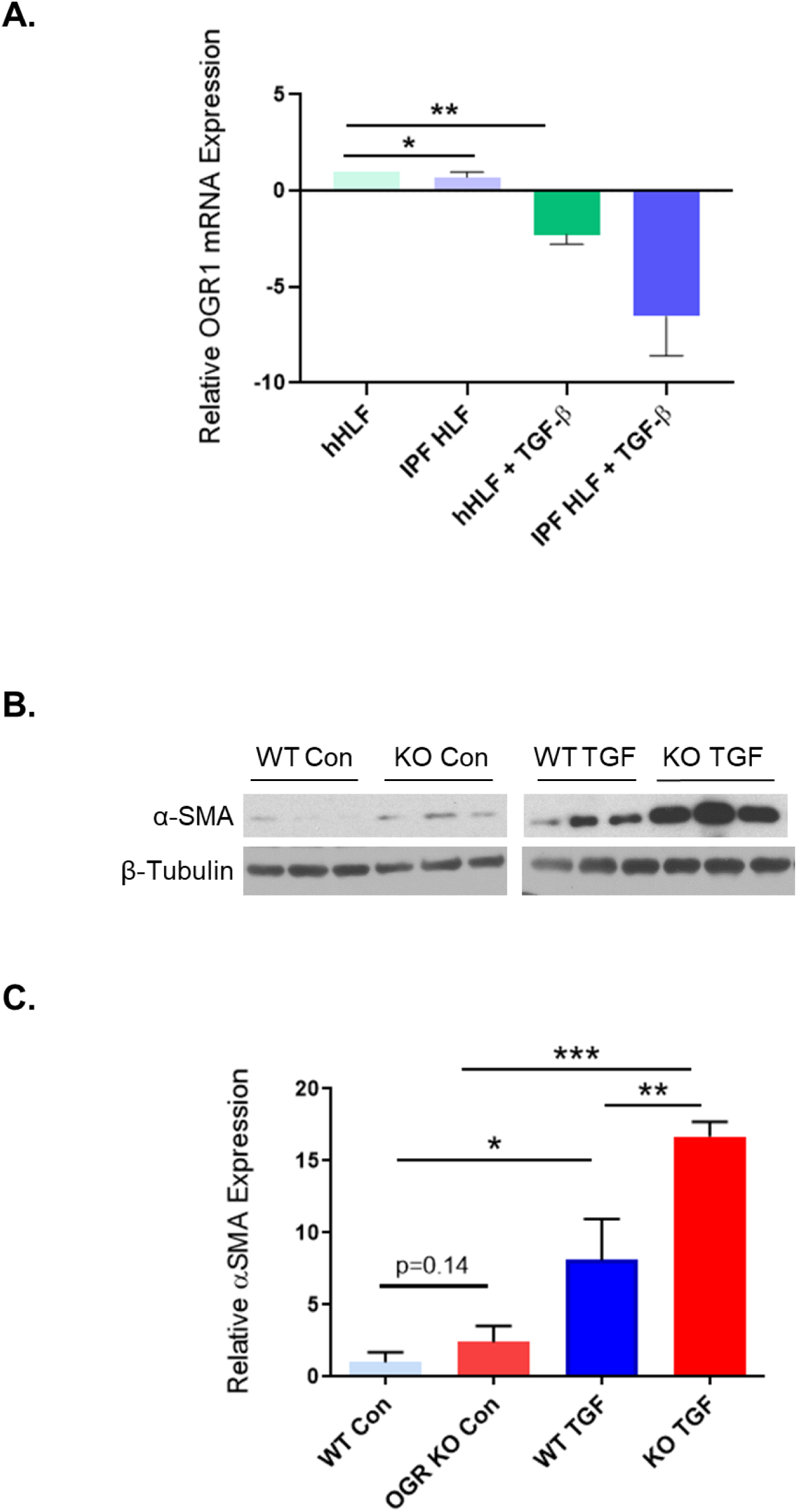
(A) Fibroblasts derived from healthy subjects and those with pulmonary fibrosis were cultured and treated with vehicle or TGF-β (1 ng/mL) for 48 hours. OGR-1 mRNA was then detected using qRT-PCR. Results represent triplicate experiments and values are normalized to 18 s mRNA expression. (B) WT & OGR1 KO mouse lung fibroblasts were cultured in the presence or absence of TGF-β (1 ng/mL) for 48 hours and alpha-smooth muscle actin expression was assessed by Western blot. (C) Densitometry analysis was performed and data represent mean α-SMA expression ± SEM normalized to β-tubulin (n=3/treatment group, repeated in triplicate). * p = 0.0034, ** p = 0.0010, *** p = <0.0001.

### OGR Overexpression Inhibits Myofibroblast Differentiation

To further assess the role of OGR1 in the negative regulation of TGF-β, we over-expressed OGR1 in wild type and OGR1 knock out MLFs and then examined changes in expression of α-SMA. Consistent with prior data, OGR1 KO MLFs demonstrated increased levels of α-SMA in the unstimulated state compared with wild type MLFs (Figure 4 A, B). OGR1 over-expression had no effect on α-SMA in wild type MLFs. However, there was a significant reduction in α-SMA expression after OGR1 was re-expressed in OGR KO cells. To assess alternative markers of myofibroblast differentiation in healthy human fibroblasts, we next over-expressed OGR1 in healthy HLFs in the presence of TGF-β and assessed changes in expression of collagen 1A1 and 3A1. In control conditions, over-expression of OGR1 had no effect on collagen expression (Figure 4 C, D). As expected, stimulation with TGF-β caused a significant increase in collagen 1A1 and 3A1 gene expression. However, when OGR1 was over-expressed, there was an approximate 50 percent reduction in TGF-β induced collagen 1A1 and 3A1 gene expression (Figure 4 C, D). These results demonstrate that the presence of OGR1 mitigates the myofibroblast differentiation stimulated by TGF-β.

**Figure 4.**
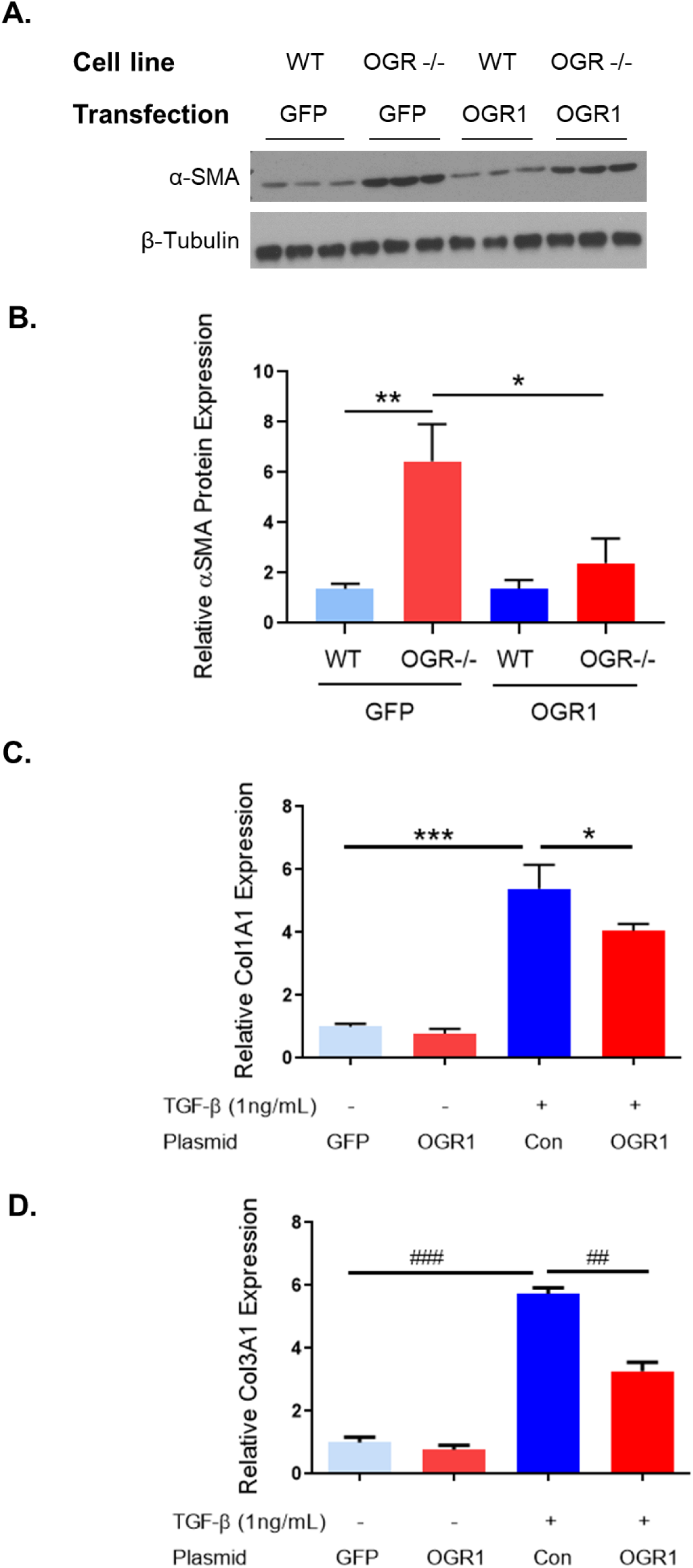
(A) Wild type or OGR1 KO mouse lung fibroblasts were treated with a plasmid expressing OGR1 or GFP for 48 hours. Protein was then harvested and Western blot was performed. (B) Densitometry analysis was performed and data represent mean α-SMA expression ± SEM, normalized to β-tubulin (n = 9 per group); * p = 0.0457, ** p = 0.0026. (C-D) Human lung fibroblasts were cultured and treated with a plasmid expressing OGR1 or GFP control. Fibroblasts were then cultured in the presence or absence of TGF-β (1ng/mL) for 24 hours. mRNA was harvested and qRT-PCR was performed. Data represent mean qRT-PCR concentration of candidate genes ± SEM relative to 18 S (n=3/treatment group, repeated in triplicate). * p = 0.0435, *** p = 0.0006, ## p = 0.0002, ### p < 0.0001.

### OGR1 Regulates Myofibroblast Differentiation via Phosphorylation of FAK

Myofibroblast differentiation has been shown to be dependent on phosphorylation of tyrosine^397^ of focal adhesion kinase (FAK) [46-48]. It has also been demonstrated that FAK integrates signals from pro-fibrotic cytokines like TGF-β[48, 49]. Therefore, we hypothesized that OGR1 may inhibit TGF-β mediated signaling by decreasing FAK phosphorylation. Indeed, FAK phosphorylation is increased by TGF-β administration and this is further enhanced by OGR1 siRNA (Figure 5 A, C). However, we did not see any effect on phosphorylation of FAK at Y^407^ (data not shown), a site that has been demonstrated to negatively regulate FAK activity[50]. In contrast, when OGR1 was over-expressed, FAK phosphorylation was attenuated (Figure 6). These findings demonstrate that OGR1 inhibits TGF-β signaling via decreased FAK auto-phosphorylation at tyrosine^397^.

**Figure 5.**
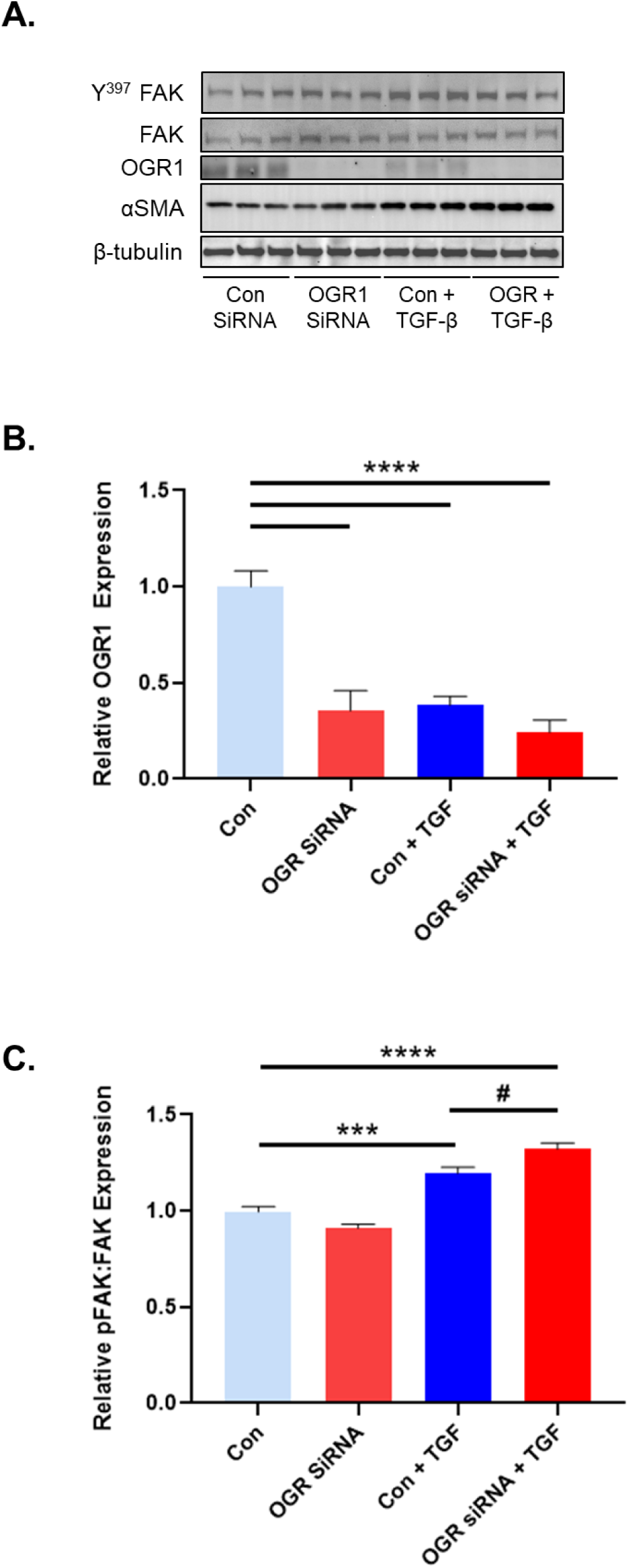
In healthy human lung fibroblasts, TGF-β significantly stimulated phosphorylation of FAK at tyrosine^397^. This phosphorylation was enhanced with OGR1 knockdown via targeted siRNA. (A) Representative western blot demonstrating effective knock down of OGR1 expression and an increase in FAK phosphorylation. (B) Densitometric analysis from multiple western blots is shown. Results demonstrate that OGR1 expression is significantly decreased in the presence of a targeting siRNA compared with a non-targeting siRNA. **** p < 0.0001. (C) Denisitometric analysis demonstrating that TGF-β significantly increases tyrosine 397 phosphorylation of FAK relative to total FAK expression; ***p = 0.0002. FAK phosphorylation at this site is enhanced when OGR1 expression is reduced with siRNA; **** p < 0.0001, # p = 0.0134.

**Figure 6.**
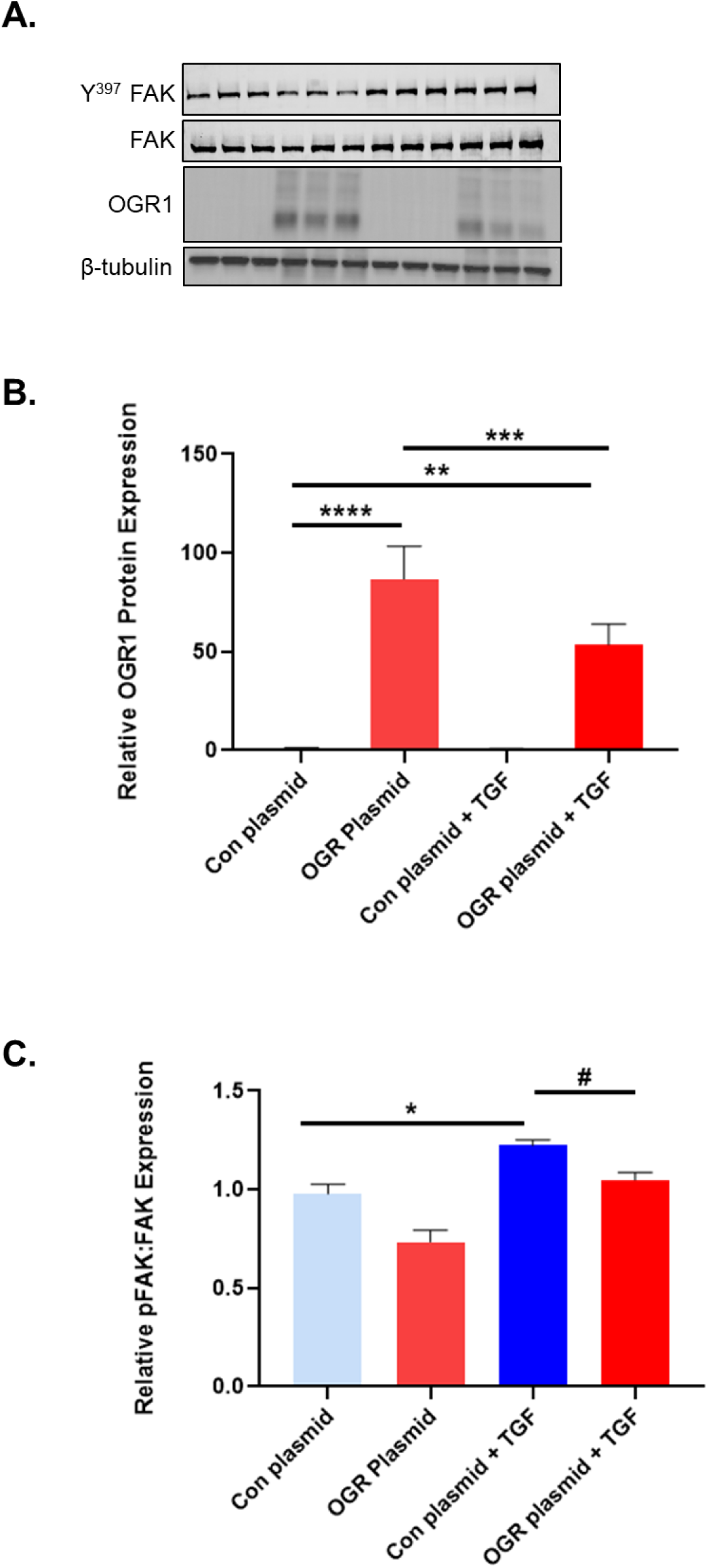
As previously demonstrated, TGF-β stimulated phosphorylation of FAK at tyrosine^397^. However, FAK activity was mitigated with OGR1 over-expression. (A) Representative western blot demonstrating an effective increase of OGR1 expression and reduction in FAK phosphorylation. (B) Densitometric analysis from multiple western blots is shown. Results demonstrate that OGR1 expression is significantly increased in the presence of an OGR1 plasmid compared with a GFP control plasmid. **** p < 0.0001, **p = 0.0033. Interestingly, TGF-β was able to significantly reduce OGR1 over-expression, *** p = 0.0005 (C) Denisitometric analysis demonstrating that TGF-β significantly increases tyrosine 397 phosphorylation of FAK relative to total FAK expression; ***p = 0.0103. However, FAK phosphorylation at this site is reduced in the presence of TGF-β when OGR1 is over-expressed; # p = 0.0203.

## Discussion

Chronic fibrotic pulmonary diseases convey a high mortality and several co-morbidities[1, 51]. An individual’s response to the available treatments varies considerably[52]. Therefore more in depth understanding of the pathogenesis and thus new therapeutic targets in pulmonary fibrosis are desperately needed. The pathopneumonic pathologic hallmark of fibrosing lung disease is the presence of fibroblast foci, sites that are enriched with activated myofibroblasts which deposit excessive extracellular matrix and disrupt normal lung architecture[7]. Myofibroblasts have a hybrid phenotype between smooth muscle cells and fibroblasts demonstrated by their ability to express contractile proteins like α-smooth muscle actin[48]. This characteristic allows myofibroblasts to repair wounds, but for reasons that are not entirely clear, pulmonary fibrosis occurs when this repair process fails to shut off. Transforming growth factor-β1 has been demonstrated to promote pathologic fibrosis in many disease states[53]. In particular, TGF-β1 has been shown to induce myofibroblast differentiation[11]. We have previously demonstrated that TGF-β leads to acidification of the extracellular space[18, 45]. This leads to increased LDH expression and activity, which promotes activation of latent TGF-β1 and creates a feed-forward pro-fibrotic signaling loop. TGF-β1 is also activated by integrins[54] and coordinates signaling from the extracellular matrix and receptors through proteins like FAK[47]. In addition, FAK expression and activity is upregulated in fibrotic foci and inactivation of FAK attenuated pulmonary fibrosis in a bleomycin mouse model[47]. In this manuscript, we sought to determine additional mechanisms by which changes in the extracellular pH are conveyed to the cell and how these signals may attenuate the pro-fibrotic effects of TGF-β1.

Here we expand the knowledge of how proton-sensing GPCRs, in particular the Ovarian Cancer G-Protein Coupled Receptor 1, negatively regulate pathologic fibrotic signaling. There are several reports that demonstrate the spatial and temporal variability of OGR1 signaling and its ramifications[27, 37-40, 55, 56]. For example, Matsuzaki et al. found that OGR1 increased connective tissue growth factor (CTGF) in airway smooth muscle cells in response to extracellular acidification. However, in a human fibroblast cell line (HFL-1) and a bronchial epithelial cell line (BEAS-2B), which both express OGR1 at higher levels than other members of this GPCR family, CTGF expression was unchanged in response to extracellular acidification[55]. This highlights the complexity of OGR1 signaling that occurs in different cells types. We found that OGR1 expression was significantly decreased in lung tissue isolated from people with IPF (Figure 1). We demonstrated that TGF-β1 downregulates OGR1 expression (Figure 3A) and that decreased OGR1 expression promotes myofibroblast differentiation (Figures 2 and 3 B,C). Finally, reconstitution of OGR1 expression or over-expressing OGR1 mitigates myofibroblast differentiation (Figure 4). These data demonstrate that OGR1 negatively regulates TGF-β pro-fibrotic signaling. We also propose that the ability of TGF-β1 to downregulate a counter-regulatory protein like OGR1 further enhances a pro-fibrotic feed forward loop.

We next illustrate that one mechanism by which OGR1 negatively regulates TGF-β1 involves inhibition of FAK phosphorylation. This pathway is even more plausible when one considers that OGR1 expression is downregulated in fibrotic foci and FAK expression/activity is significantly upregulated in the same areas[47]. Activation of FAK is largely dependent upon phosphorylation of several tyrosine residues, including Tyr^397^, Tyr^576^, Tyr^925^, among others. Phosphorylation at these residues stimulates several downstream kinases including phosphatidylinositol 3-kinase (PI3K), p130CAS, Ras-GEF, Shc, ERK, and Grb2[50, 57]. Though the specific mechanism by which OGR1 affects FAK phosphorylation deserves additional investigation, our data demonstrate that OGR1 antagonizes TGF-β-mediated FAK activity.

It is interesting to note that the OGR1 global knock out mouse does not develop an overt phenotype. There is a growing recognition that genetic variations play a role in the development and clinical course in idiopathic pulmonary fibrosis. However, only about 1/3 of IPF diagnoses are attributed to common genetic variants[58]. Therefore, at our present level of understanding, it does not appear that the majority of IPF cases can be explained by genetic alterations alone. We plan to further characterize the response of OGR1 knock out mice to bleomycin challenge. We anticipate that they will be more susceptible to bleomycin injury. The other common theme in clinical IPF trials is that aiming at one target may not be enough to demonstrate a significant clinically meaningful outcome due to pro-fibrotic signal redundancy. Indeed, the current FDA approved anti-fibrotic medications slow the progression of disease but patient-reported symptoms were largely unchanged. Pooled data from the INPULSIS and TOMORROW trials involving nintedanib demonstrated a reduction in acute exacerbations[59] and secondary endpoint analysis with pirfenidone and nintedanib suggest an improvement in all-cause and IPF related mortality[60, 61]. These additional benefits need to be confirmed with larger randomized clinical trials. Given that the absence of OGR1 does not confer an overt pulmonary fibrotic phenotype, we recognize that solely targeting OGR1 as a new anti-fibrotic strategy is misguided. Rather, we suggest using a multimodal approach to IPF to combat the redundant nature of pro-fibrotic signaling pathways. We propose that the preservation of OGR1 expression and activity is an important addition to current and yet to be approved medications used to treat pulmonary fibrosis.

## Acknowledgements

The authors would like to acknowledge Wade Narrow and Jennifer Judge, PhD for technical assistance.

